# deStruct: Accurate Rearrangement Detection using Breakpoint Specific Realignment

**DOI:** 10.1101/117523

**Authors:** Andrew McPherson, Sohrab Shah, S. Cenk Sahinalp

## Abstract

We propose that a breakpoint specific alignment procedure would improve breakpoint prediction. Our method, deStruct, uses multiple stages of realignment and clustering to progressively refine breakpoint prediction quality and accuracy. We show using simulated data that deStruct predicts breakpoints with higher sensitivity and specificity than existing breakpoint prediction tools.

## 1 Introduction

Improvements to high-throughput whole genome sequencing technologies have resulted in increased read length with no corresponding increase in fragment length. As a result, reads produced from fragments that span breakpoints are more likely to interrupt read sequences, complicating alignment and breakpoint prediction. The primary goal of general read alignment tools [7, 8, 5] is accurate alignment of concordant reads. Many existing breakpoint discovery methods then search for evidence in the alignments produced by a general read alignment tool [6, 11, 1]. We propose that a breakpoint specific realignment strategy may increase accuracy for breakpoint prediction.

The deStruct breakpoint prediction method uses a series of realignment and clustering steps to progressively refine breakpoint prediction quality and accuracy. In brief, we first assume that a significant number of reads supporting real breakpoints will be split by those breakpoints. As we do not know, a-priori, the location of the split in each read, the first step will be to identify the optimal prefix alignment of each read. Pairs of read prefix alignments with low alignment probability are filtered, with split read alignment attempted based on the remaining prefix alignments. Read prefix alignments are clustered, and each read in a cluster is realigned to the breakpoints nominated by the top scoring split read alignments in that cluster. The final output of the realignment and clustering step is a set of paired end read alignment clusters, and alignment likelihoods per read per cluster. We then solve an optimization problem to select the most likely set of clusters, and the most likely set of read to cluster assignments. Below we detail the overall objective of the method followed by the specifics of each step.

## 2 Method

### 2.1 Objective

The objective of deStruct is to identify breakpoints, and assign read alignments to those breakpoints. Herein we describe the specific objective we attempt to (approximately) maximize with the deStruct method. Let *b_k_* be a binary indicator that breakpoint *k* exists, and let *z_i_∈*{1..*K*} be a per read indicator that read *i* is assigned to breakpoint *k*. Let *r_i_* represent the observed sequence of read *i*, where 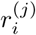 represents end *j* of read *i*.

Calculate a prior over all *B* as given by Equation 1, where *P*(*b_k_* = 1) is assumed small as any randomly selected breakpoint in the genome is assumed rare.

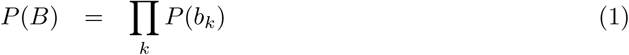

Calculate the probability of read to breakpoint assignments as given by Equation 2, where the probability of assigning a read to a breakpoint *k* is fixed at 0 if *b_k_* indicates the breakpoint does not exist ( Equation 3).

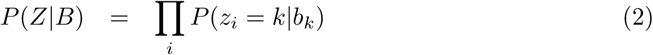

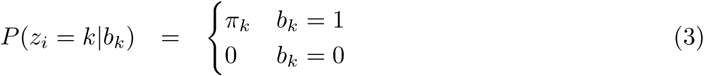

We approximate the probability of observing the sequence of read *i* given its assignment to breakpoint *k* ( Equation 4) as the argmax over possible alignments *a* ( Equation 5), a more tractable calculation that marginalizing over *a*.

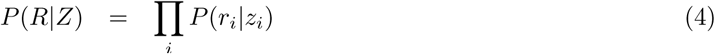

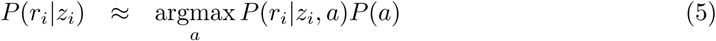

Finally, model the likelihood of read i as a product of the probability of the inferred template length tl(*a, k*) and the alignment score 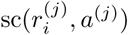 of each end ( Equation 6).

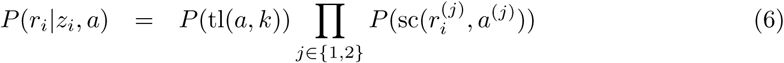

Our objective is then to infer the *Z* and *B* that maximize the likelihood given by Equation 7.

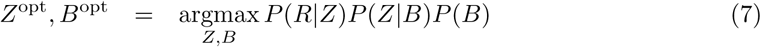

### 2.2 Algorithm overview

Discordant read alignment uses a seed and extend strategy to independently align discordant read pairs to the reference genome. Read alignments, including partial alignments, are filtered according to mapping specificity as given by an approximate calculation of the alignment posterior (Section 2.3.1). An optimal split alignment is calculated for each remaining alignment pair, allowing for insertions at the breakpoint, and normalized to remove ambiguity resulting from sequence homology at the breakpoint (Section 2.3.2).

Paired alignments are clustered according to the likelihood they originate from the same unknown breakpoint (Section 2.4). For each cluster, a set of potential breakpoints are selected, including breakpoints supported by high numbers of split alignments and breakpoints constructed from partial alignments. Reads are realigned to potential breakpoints of associated clusters to provide a more accurate alignment score sc(*r^(j)^, a^(j)^*).

The result of the alignment and clustering steps are a set of potential breakpoints, potential assignments of reads to breakpoints, and a likelihood for each assignment. Our choice of procedure for inference is motivated by the belief that *P(b_k_)* is infinitesimally small, and dominates the likelihood given by Equation 7. Thus, the first step in our optimization procedure is to identify the mapping of reads to breakpoints that minimizes the number of events. Formulated as an instance of setcover, this subproblem can be solved approximately with the well known greedy approximation algorithm [4]. Finally, each read is assigned to the position and breakpoint that maximizes the likelihood ( Equation 4) of that read.

To summarize, the algorithm involves the following steps:

1. breakpoint specific alignment to produce spanning and split alignments
2. clustering of spanning alignments, enumerating potential breakpoints, and realignment of reads to potential breakpoints
3. assignment of reads to breakpoints to minimize number of breakpoints
4. assignment of reads to positions/breakpoints to maximize likelihood

More detail is provided in the following sections.

### 2.3 Breakpoint specific alignment of discordant reads

**Figure 1:**
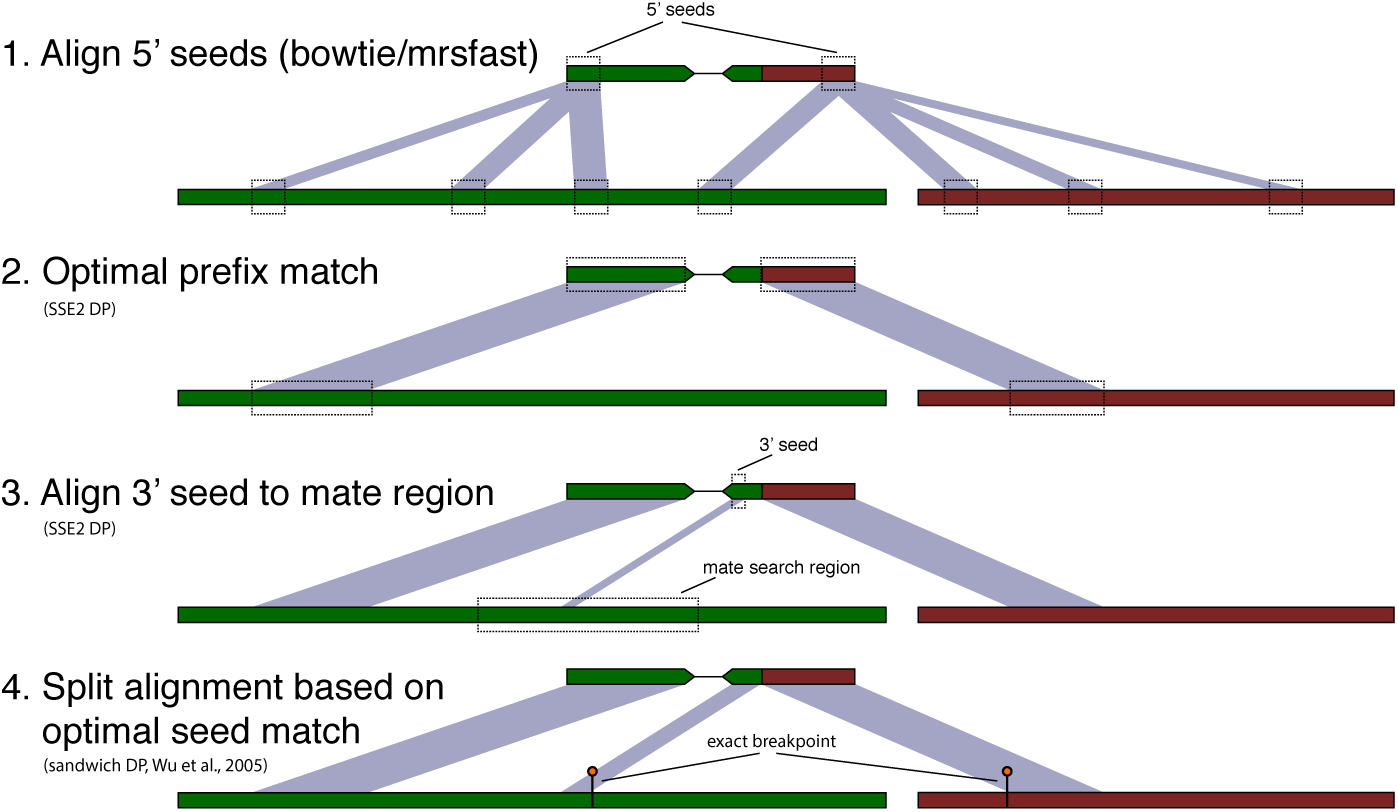
Breakpoint specific read alignment. 1) 5-prime seeds are aligned to the genome to obtain the top n alignments within a stratum of alignment scores. 2) 5-prime seed alignments are extended using SSE2 optimized banded dynamic programming. 3) 16 nt 3-prime mate seeds are aligned within m nt of the 5-prime seed matches where m depends on the length of the fragments. 4) 3-prime mate seed alignments are extended and combined witih 5-prime mate alignments to identify an optimal split.

#### 2.3.1 Probabilistic partial alignment for identification of potential split reads

We propose a seed and extend alignment strategy. For each read, a comprehensive set of alignment positions is identified for 5-prime seeds (first *k* nt of each read). Seed alignments are extended to a length that optimizes an alignment score (with linear gap penalty). Extension uses SSE2 optimized dynamic programming (DP) to calculate DP matrix for the alignment, to be potentially used in the subsequent split alignment step. For reads with extensions of different optimal lengths, the score for each extension is taken at the maximum of the optimal lengths, minus *q* nucleotides to account for spurious matches. The resulting partial alignments are filtered according to their specificity as given by an alignment posterior, in addition to the posterior probability the read is in fact concordant.

We adopt a strategy for calculation of alignment posteriors similar to [12]. Let *x*^(1)^, *x*^(2)^ be alignment locations of paired ends *r*^(1)^, *r*^(2)^ respectively. Let *C* be the set of alignment location pairs that are concordant, and let *d* be the prior probability that the true alignment locations are discordant. Calculate the prior probability of observing alignment pair *x*^(1)^, *x*^(2)^ as given by Equation 8. Note that [12] use a prior that assumes pairs of discordant alignments are drawn independently from the genome length, resulting in an infinitesimally small prior for all discordant alignments. By contrast, we assume that discordant status and one alignment location in the pair is sufficient to fully specify a prior on the pair, since a true tumour genome will not contain all possible pairs of positions as breakpoints. The posterior probability of one location of a pair is calculated as given by Equation 9. The numerator *N* and denominator *Z* can be calculated efficiently as given by Equations

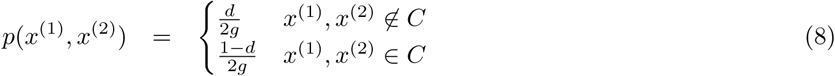

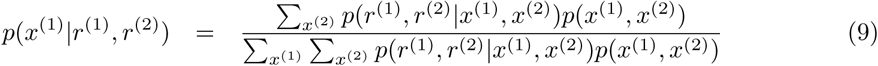

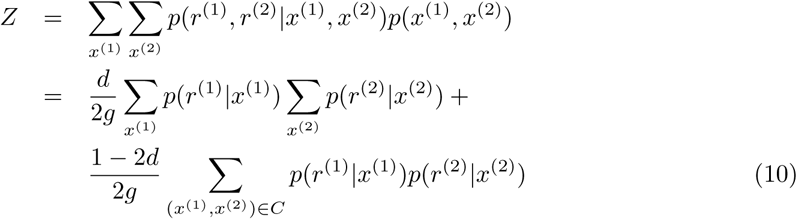

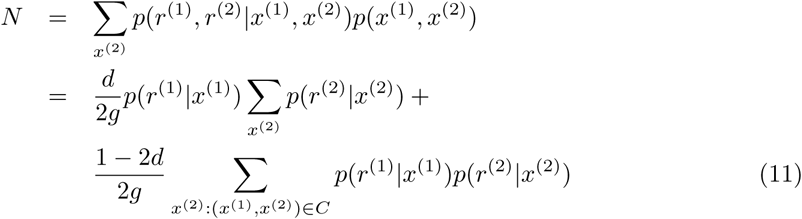

Also calculate the probability that any of the concordant alignments of *r*^(1)^ and *r*^(2)^ are true Equation 12). Alignments exceeding a threshold probability of concordance are filtered.

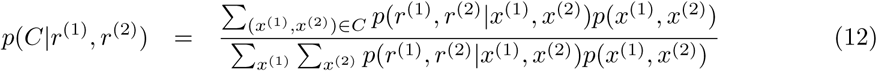

We model the probability *p*(*r*^(1)^|*x*^(1)^) as the likelihood of observing read *r*^(1)^ with origin *x*^(1)^ in the genome (*p*(*r*^(2)^|*x*^(2)^) is defined similarly). Let sc(*r*^(1)^,*x*^(1)^) denote the score for the optimal alignment of *r*^(1)^ to *x*^(1)^. We calculate *p*(*r*^(1)^|*x*^(1)^) ≈*p*(sc(*r*^(1)^, *x*^(1)^)) using the distribution of alignment scores of concordantly aligning reads. In brief, we align a subset of the reads to the genome using an aligner configured for maximum sensitivity. We then calculate the distribution of alignment scores for all concordant reads and use this density as an estimate for *p*(sc(*r*^(1)^, *x*^(1)^)). We calculate *p*(sc(*r*^(1)^, *x*^(1)^)) for all substrings of *x*^(1)^ to allow us to calculate *p*(*r*^(1)^|*x*^(1)^) for partial alignments of any length.

#### 2.3.2 SSE2 optimized split alignment

Filtered partial alignments obtained as specified in Section 2.3.1 represent the starting point for split read alignment and subsequent identification of breakpoints. For a read whose mate is split by a breakpoint, the 3-prime end of the mate should align within *m* nucleotides of the read where *m* is the length of the sequenced DNA fragment. For each read alignment, we calculate the optimal alignment of a 3-prime seed of the reads mate to a window adjacent to the read alignment. For efficiency, the 3-prime seed is taken to be the last 16 nt of the mate read, since this is the size of a single SSE2 register. Alignment uses dynamic programming implemented with SSE2 instructions. Seed matches that exceed a threshold are extended similarly to 5-prime seed matches to produce a DP matrix for the alignment.

An optimal split alignment can be calculated efficiently from two DP matrices: one created for alignment to the 3-prime seed match location, and one created for the (reverse) alignment to the 5-prime seed match location. Let A_1_[*i*, *j*] and A_2_[*i*, *j*] be these two DP matrices respectively, with rows corresponding to read sequence positions and columns to locus positions. Let *I* be the length of the read sequence and *q* the linear penalty for insertion at the breakpoint. Our objective is to find *i*_1_, *i*_2_, *j*_1_, *j*_2_ such that *i*_1_+*i*_2_ ≤ *I* to maximize SPLIT_OBJ defined in Equation 13. Note that the requirement that *i*_1_+*i*_2_ ≤ *I* is required since we assume that each segment of the read has a single origin: locus 1, locus 2, or novel inserted sequence.

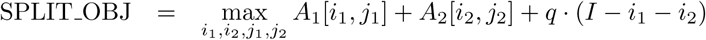

Let *B*_1_[*i*] and *B*_2_[*i*] be the max score for each read sequence length aligned to locus 1 and 2 respectively. Let *C*_1_ [*i*] be the max score including potential insertion for the read sequence aligned to locus 1. Let *CX*_1_[*i*] be the last aligned read position before introduction of inserted nucleotides that produces maximum scores for *C*_1_[*i*].

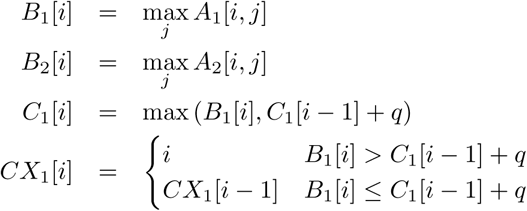

We can calculate an optimal *i*_i_, *i*_2_ by maximizing *C*_1_[*i*] +*B*_2_[*I*−*i*].

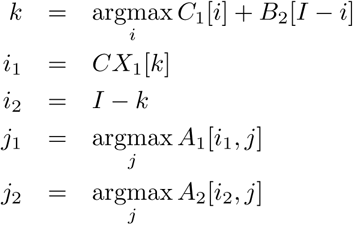

The solution *i*_1_, *i*_2_, *j*_1_, *j*_2_ is not guaranteed to be unique. Of particular interest is ambiguity due to homologous sequence at the breakpoint. Homologous breakpoint sequences will produce solutions of the form {(*i*_1_, *i*_2_, *j*_1_, *j*_2_), (*i*_1_+1, *i*_2_− 1, *j*_1_ + 1, *j*_2_− 1),…, (*i*_1_+*h*, *i*_2_ − *h*, *j*_1_+*h*, *j*_2_ − *h*)} with *i*_1_+*i*_2_ = *I*, and where *h* is the number of homologous nucleotides. To unambiguously represent a breakpoint predicted by such an alignment, we select the breakpoint positions *j*_1_ and *j*_2_ such that *j*_1_ < *j*_2_ and *j*_1_ is minimal. Thus, the triple (*j*_1_, *j*_2_,*h*) represents breakpoints (*j*_1_, *j*_2_) and (*j_1_ + h*, *j*_2_ − *h*) but not (*j*_1_ − 1, *j*_2_+ 1) or (*j*_1_+*h*+ 1, *j*_2_ − *h* − 1)

### 2.4 Clustering paired alignments

Let 𝒜 be the set of all paired end alignments of a paired end read library (or libraries). The objective of the clustering process is to determine, from 𝒜, the most likely set of breakpoints and the most likely assignment of alignments in 𝒜 to those breakpoints. We cluster 𝒜 using a two step process. We first perform a minimum spanning tree based clustering step to quickly separate the data, followed by a more refined clustering step using a mixture model.

Consider a paired end alignment 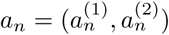. Let 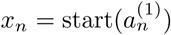 and 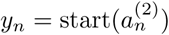 be the positions in the genome to which the first nucleotide of each read aligns. Let *μ_n_* and *σ_n_*be estimates of the mean and standard deviation for fragment *n*. The values of *μ_n_* and *σ_n_* are estimated per input sequencing dataset, allowing our clustering algorithm to be used to cluster reads from multiple datasets simultaneously.

#### 2.4.1 Minimum Spanning Tree Clustering

We begin by applying a transformation to the paired end alignment data that will desegregate paired end alignments resulting from the same breakpoint. Let *B* = *(s,t)* be a breakpoint joining genomic positions *s* and *t*. A set of paired end alignments resulting from *B* and having similar *μ_n_* will have *x_n_* and *y_n_* values near the diagonal line *s* − *x_n_* + t − *y_n_ − μ_n_* = 0. Paired end alignments with different *μ_n_* will segregate each on their own line defined similarly (Figure ??). Thus we apply a transform to ‘subtract out’*μ_n_* such that the transformed *x_n_*, *y_n_* all lie on the same diagonal line. An additional scale ensures that the transformed positions *v_n_, w_n_* from the same breakpoint will lie approximately within a unit square. The full transformation is given by Equation 15. We use *q* = 3, since 99% of the fragments will have a length within 3 standard deviations of the mean for those fragments. The increased separation between clusters is illustrated in Figure ??.

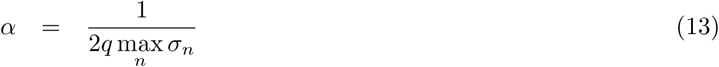

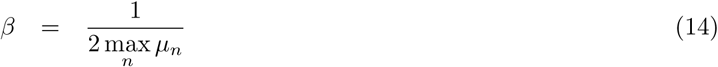

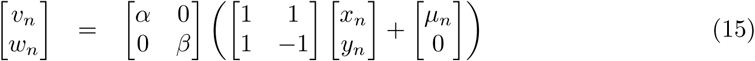

**Figure 2:**
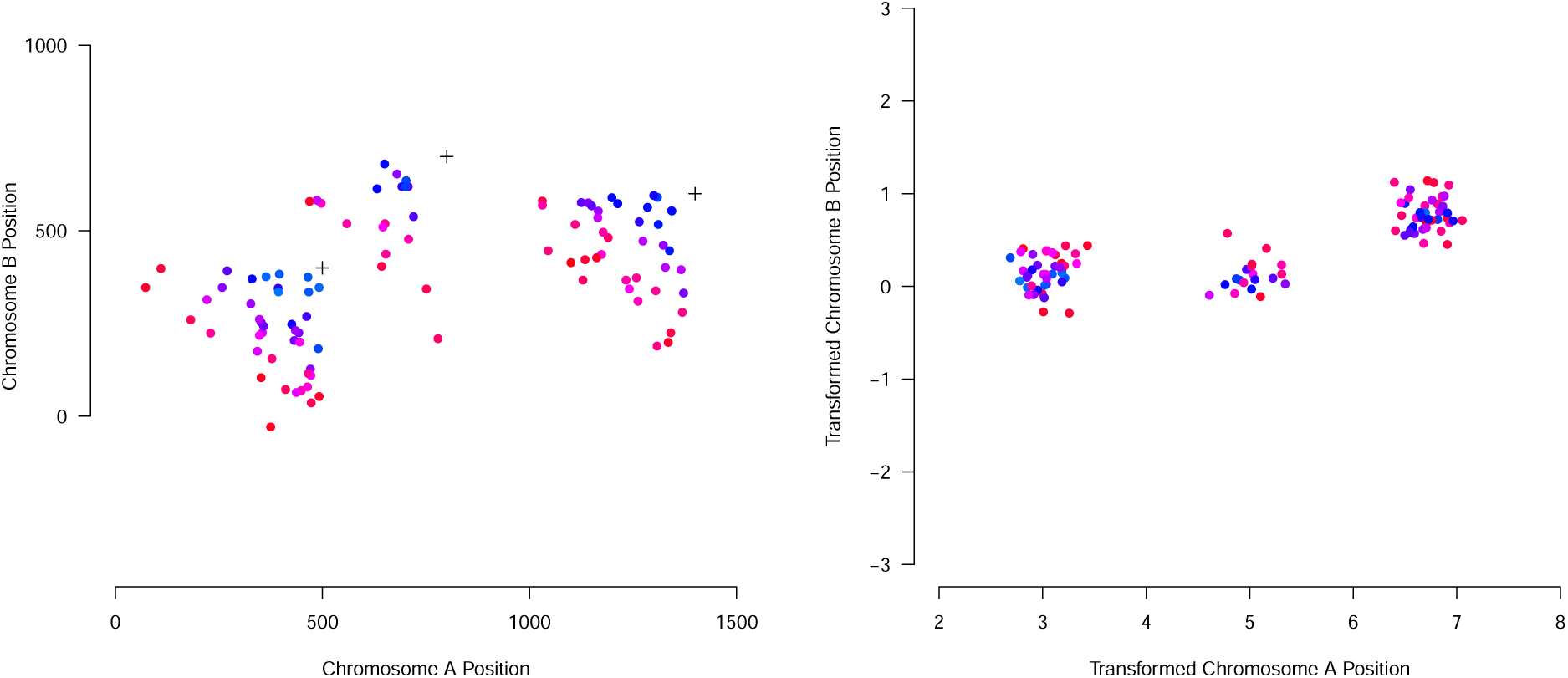
Transformation of simulated paired end alignments. Points are coloured according to *μ_n_*⋅ (a) Positions of mate pairs derived from 3 simulated breakpoints, showing the tendency of alignments with similar *μ_n_* to segregate along lines given by *s − x_n_*+*t*−*y_n_ −μ_n_* = 0. (b) Transformation of mate pair positions showing increased separation between clusters.

Next we apply a minimum spanning tree based clustering to the *v_n_* and *w_n_* values to provide an initial clustering. To do this we first calculate the Delaunay triangulation of all (*v_n_*, *w_n_*) points, which can be done efficiently in 𝒪(*n* log *n*) time. Next we cut all edges in the Delaunay triangulation with length greater than *h*, and take the connected components of the resulting graph as the initial clusters. The euclidean minimum spanning tree for a set of points in the 2D plane is a subgraph of every Delaunay triangulation of those points. Thus points (*v_n_*, *w_n_*) and (*v_m_*, *w_m_*) are in different initial cluster only if the euclidean distance between (*v_n_*, *w_n_*) and (*v_m_*, *w_m_*) is greater than *h*. We use *h* = 2.

#### 2.4.2 Mixture Model Clustering

Each initial cluster produced by the previous step is refined into a set of smaller clusters using collapsed gibbs sampling of a dirichlet process mixture model. Given a breakpoint formed by joining the sequence to the left (upstream) of position s in chromosome *X* to the sequence to the left (upstream) of position *t* in chromosome *Y*, we can write the likelihood of alignment *A* = (*x*, *y*) given breakpoint *B* = (*s*, *t*) as given by Equation 16.

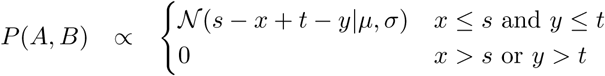

Note that for breakpoints that involve sequence downstream of position *s* and or *t*, Equation 16 can be used by reversing the chromosome sequence and associated alignments, which is mathematically equivalent to negating the alignment and break-end positions.

Given a set of read alignments, we would like to assign each alignment to a single cluster so as to maximize the likelihood. As we don’t know a-priori the number of required clusters, we use a Dirichlet Process mixture model to learn the number of clusters from the data. The Dirichlet Process prior can be written as given by Equation 16 with concentration parameter ζ.

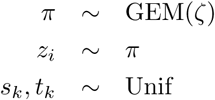

We used collapsed Gibbs sampling to efficiently sample from the posterior over read alignment clusters, or equivalently, breakpoints. Specifically, we use ‘’algorithm 3” from [10], as given by Equation 16. Let **A**_*k*_ be the alignments in cluster *k*, and let *A_n_* be an unassigned alignment. At each step in the sampling process, alignment *n* is removed from the cluster to which it is assigned and reassigned at random according to a probability calculated by Equation 16.

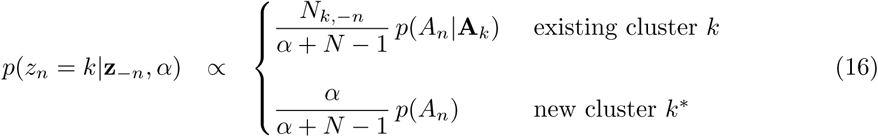

Collapsed Gibbs requires efficient calculation of the predictive probability of a cluster *p*(*A_n_*|**A**_*k*_), evaluated at the new datapoint *A_n_*. Fortunately, the likelihood given by Equation 16 contains symmetry that can be leveraged for producing a near closed form solution for the marginal probability of a cluster of alignments. First consider the case of calculating *P*(*A*), the marginal probability of a single alignment, required for normalizing *P*(*A* | *B*). By inspection *P*(*A*) is in fact independent of the value of *A* = (*x*, *y*), thus we can assume *x* = *y* = 0. Next, we observe that the value of *P*(*A*, *B*) is equal on the diagonal line *v* = *s* + *t* for *v* constant, given *s*, *t* ≥ 0. The length of this diagonal line is 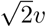 and it is 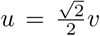 along the perpendicular diagonal. The area of a vanishingly small rectangle surrounding *v* = *s* + *t*: *s*, *t* ≥ 0 can be calculated as 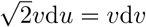, and the value of *P*(*A*) for all points in that rectangle is 𝒩(*v*|*μ*, *σ*^2^). Thus we can calculate *P*(*A*) as given by Equation 17, where we have substituted *v* = *σw* + *μ* in the third step.

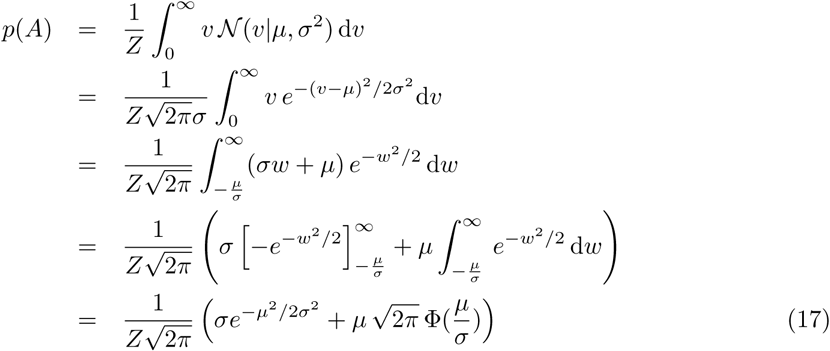

We leverage the same symmetry to calculate the marginal likelihood of a cluster of alignments. For a cluster of alignments **A**, the region of positive likelihood is *u* ≥ max_*x*_*n*_∈**A**_, *v* ≥ max_*y*_*n*_∈**A**_, thus *v* = 0 at the point *u* = max_*x*_*n*_∈**A**_ and *v* = max_*y*_*n*_∈**A**_. We will calculate *p*(**A**) as given by Equation 18.

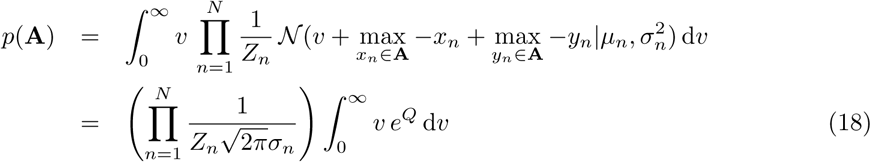

Define *w_n_* as given by Equation 19 and calculate the exponent *Q* as given by Equation 20, where we have subsumed some of the calculations based on *σ_n_* and *w_n_* into variables *α*, *β*, and

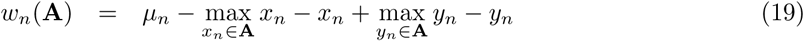

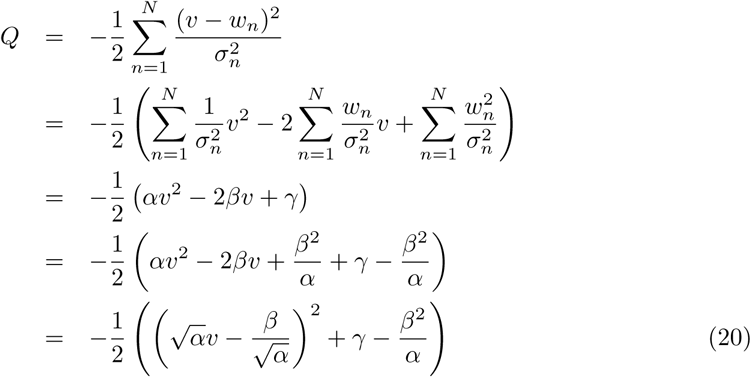

Substitute *u* for *v* with Equations 21 and 22.

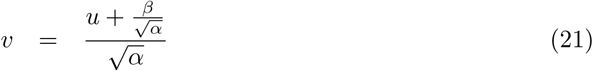

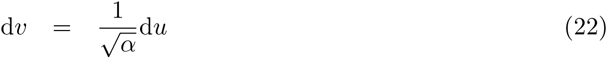

Finally, calculate *p*(**A**) in terms of on *α*, *β*, and *γ* as given by Equation 23.

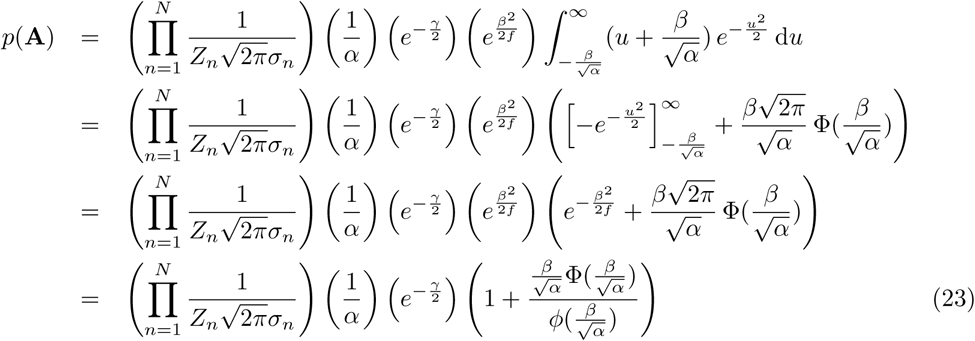

To perform collapsed Gibbs sampling efficiently, we require the ability to calculate *p*(*A_n_*|**A**_*k*_) = *p*(**A∪**{*A*'})/*p*(**A**) efficiently based on sufficient statistics of each member of **A**_*k*_. Define *r*(**A**, *A*') as given by Equation 24.

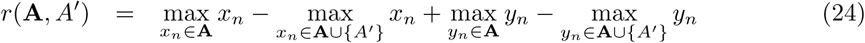

Updates to *α* are given by Equation 26.

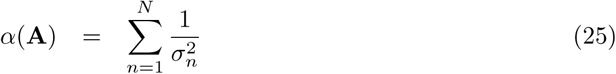

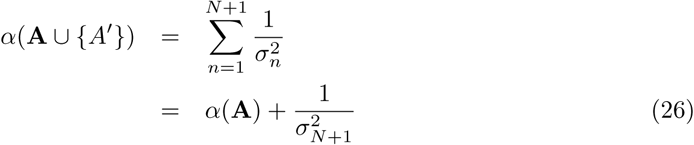

Updates to *β* are given by Equation 28.

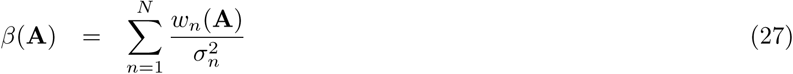

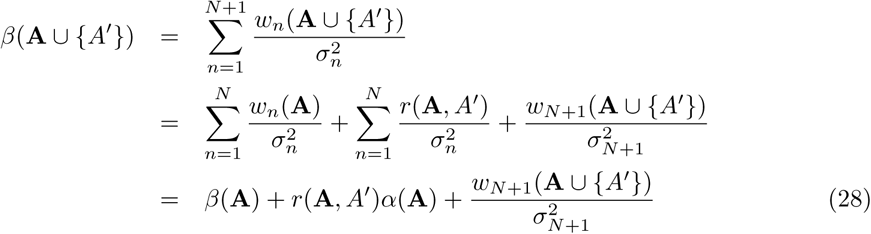

Updates to *γ* are given by Equation 30.

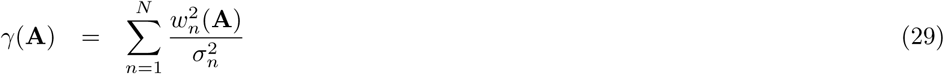

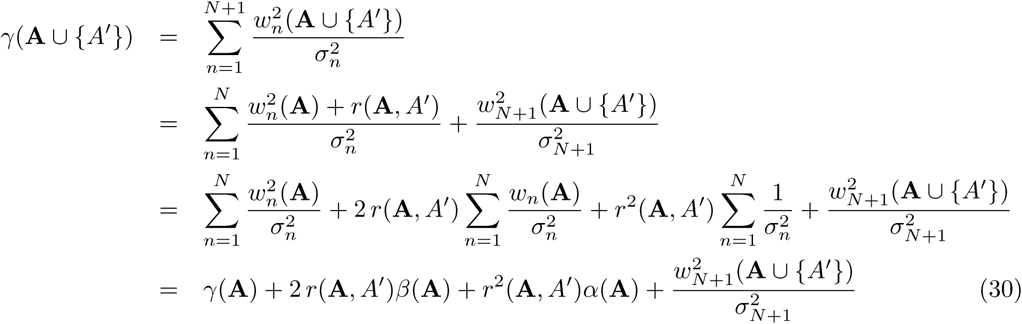

The algorithm proceeds as follows. Read pair alignments are shuffled. For each read pair alignment, remove the alignment from the cluster to which it is currently assigned if it exists, calculate Equation 16, normalize, and randomly assign the alignment to a new cluster based on the calculated probability. Calculating Equation 16 can be done in constant time by updating *α*, *β*, and *γ* and recalculating *p*(**A**) using Equation 23. Adding and removing from a cluster can be done in constant time providing read alignment positions are maintained in a priority queue, allowing for efficient updates of max_*x_*n*_*∈**A**_ and max_*y_*n*_*∈**A**_. Finally, at each step we keep track of the joint marginal probability of the full set of alignments, which can be calculated as given by Equation 31, and select the clustering that maximizes the joint marginal after a specified number of iterations.

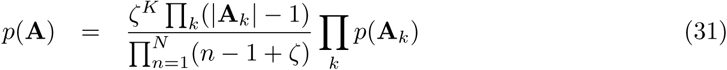

### 2.5 Selecting the optimal set of breakpoints

The optimization procedure for selecting the most likely set of real align works in two stages. First, we select the most parsimonious set of read alignment clusters, and assignments of reads to those clusters. We assume that the prior probability of any individual breakpoint is sufficiently low, that minimizing the number of alignment clusters (thus minimizing the number of breakpoints) will always maximize Equation 7. The subproblem of minimizing the number of clusters can be solved approximately in polynomial time using a reduction to an instance of setcover, as previously described[4].

We now have an assignment of read alignments to clusters, a set of breakpoints per cluster nominated by analysis of split read alignments, and a set of refined read alignment likelihoods calculated from the realignments to those breakpoints. Per read alignment cluster, we calculate a likelihood for each breakpoint at the product of individual breakpoint specific read alignment likelihoods. Finally, we take the maximum likelihood breakpoint for each cluster.

### 2.6 Identification of balanced rearrangements using breakpoint graphs

To identify balanced rearrangements, we propose the following optimization problem. We wish to identify the set of balanced rearrangement breakpoints that simultaneously maximize the number of breakpoints assigned to balanced rearrangements, and minimize the length of loss and gain edges, or a weighted combination thereof. Our solution uses a similar combinatorial formulation to that used by ReMiXT[9]. The problem can be solved by identifying the minimum cost set of edge disjoint alternating cycles (corresponding to balanced rearrangements) for a Genome Modification Graph with appropriate edge weights: ‘+’ and ‘−’ segment edges are given a positive weight scaled by length, ‘+’ breakpoint edge weights are given a negative weight, ‘−’ breakpoint edge weights are given infinite weight, telomere edges are removed and reference bond edges are given weight 0. Minimum cost perfect matching could then be used to identify all balanced rearrangement breakpoints simultaneously.

## 3 Results

### 3.1 Simulating Realistic Datasets

We used a novel approach for simulating genome sequence data to benchmark the performance of deStruct against existing breakpoint prediction software. Our approach starts with real sequencing data from a normal sample (we used HCC1395BL [3]). Similar to BamSurgeon[2], we split the normal sample into a a set of reads that will be used for the simulated normal sequencing dataset, and a set of reads that will be used for a tumour sequencing dataset. We then spike in breakpoint reads into the tumour sequencing dataset.

Spiking the tumour dataset with breakpoint reads works as follows. We first select random break-ends such that the resulting breakpoint has the desired breakpoint sequence homology. Next we create the breakpoint sequences, adding inserted sequence at the breakpoint where specified in the simulation. Finally, we simulate genome sequence reads produced by each breakpoint.

Our objective is for the breakpoint reads to be as similar to background of concordant reads as possible. Thus we use the following approach to simulate breakpoint reads. First, we pick a random location in the original genome that has sufficient read depth. Next, we virtually transplant the reads onto the breakpoint sequence template, and modify the sequence of the read to match the breakpoint sequence. Specifically, for each matched nucleotide in the original alignment of the read, we generate a match with the breakpoint sequence by modifying the nucleotide at the matched position in the read. For each mismatched nucleotide we generate a mismatch at the mismatched position in the read. We generate mismatches in a deterministic way so as to preserve consistent mismatches of the same nucleotide at the same position as would happen for single nucletotide polymorphisms. For small insertions, we preserve the inserted sequence in the read, and for small deletions, we ensure the read is remapped to the breakpoint sequence so as to preserve the deletion.

### 3.2 Benchmarking Results

We simulated 10000 breakpoints on a reference genome consisting of chromosomes 20 and 21 of hg19. For each breakpoint, we randomly selected a target homology uniformly from the range 0 to 4. For 10% of the breakpoints with homology 0, we randomly selected an inserted sequence with maximum length 5. Breakpoint sequence reads were simulated and spiked in as described above, with 25% of the original coverage in HCC1395BL. We then ran deStruct, LumpySV[6] and Delly[11] on the simulated datasets and identified correctly predicted breakpoints as those for which the break-ends were within 200nt of the simulated break-ends. For each tool, we calculated the number of true positives, and false positives for a series of thresholds on a set of features that included the number of spanning reads, the number of split reads, and for deStruct the breakpoint log likelihood. We then selected the feature/threshold combination that maximizes prediction accuracy (calculated as TP + TN/(P + N)).

Accuracy for each tool is shown in Table 1, showing deStruct with higher accuracy than either delly or lumpy. Figure 3 shows the number of breakpoints with correct and incorrect prediction of breakpoint sequence, homology, and inserted sequence, in addition to the number of breakpoint for each tool that were missed entirely. deStruct outperforms the other tools with respect to calculation of breakpoint features, primarily because it is able to recover split reads for a higher proportion of breakpoints (Figure 4).

**Figure 3:**
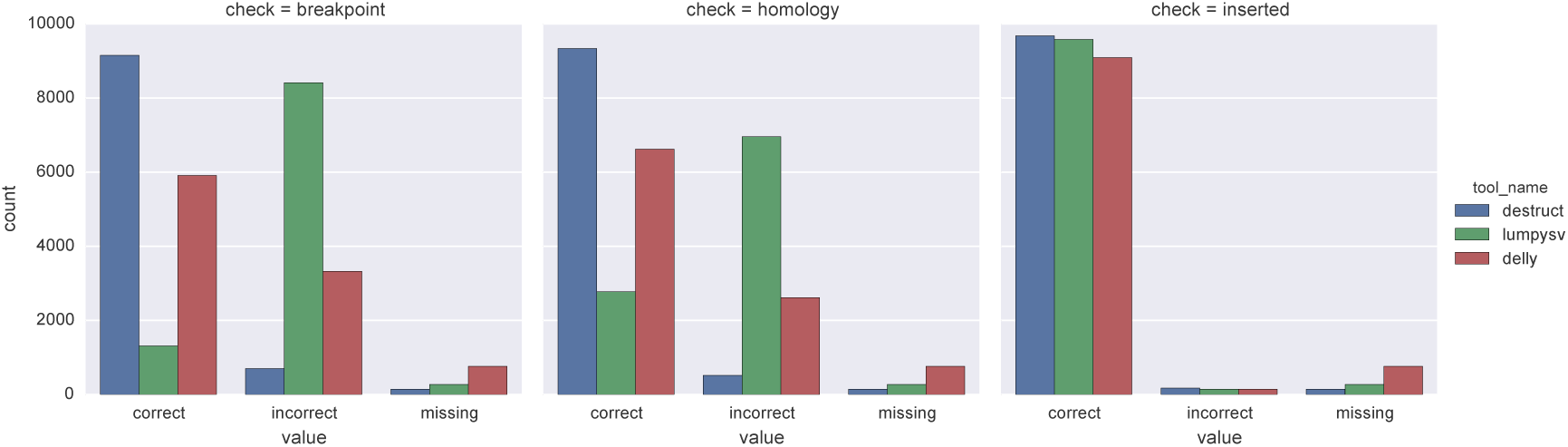
Accuracy of exact breakpoint position (left), breakpoint homology (middle), and sequence inserted at the breakpoint (right) evaluated for destruct, delly, and lumpysv based on simulated breakpoints.

**Figure 4:**
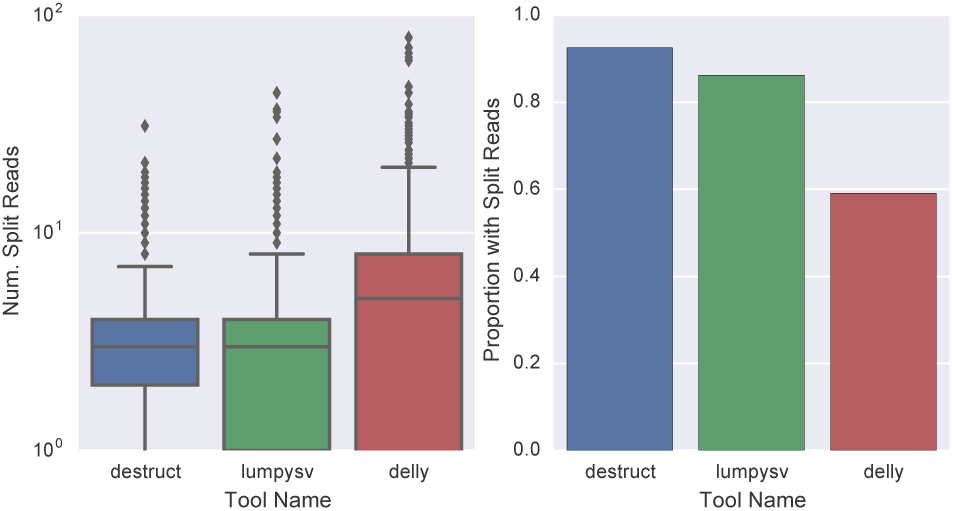
Boxplots of the log of number of split reads recovered for breakpoints for breakpoints with at least 1 split read (left) and proportion of breakpoints with at least 1 split read (right) for destruct, delly, and lumpysv based on simulated breakpoints.

**Table 1:**
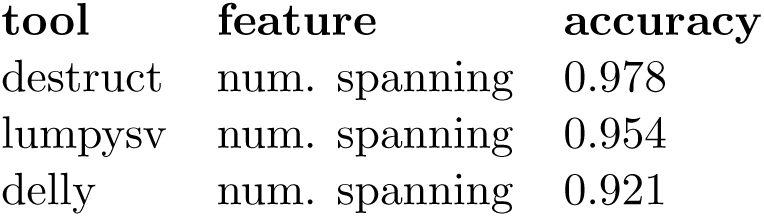
Accuracy per tool for an optimal feature/threshold combination.

Finally, Figure 5 shows as plot of the true positive and false positive counts for a series of thresholds on the best feature by accuracy for each tool. DeStruct shows considerably higher sensitivity on the simulated data, recovering a significantly higher proportion of true breakpoints. Fiowever, deStruct also predicts a higher number of false positives.

**Figure 5:**
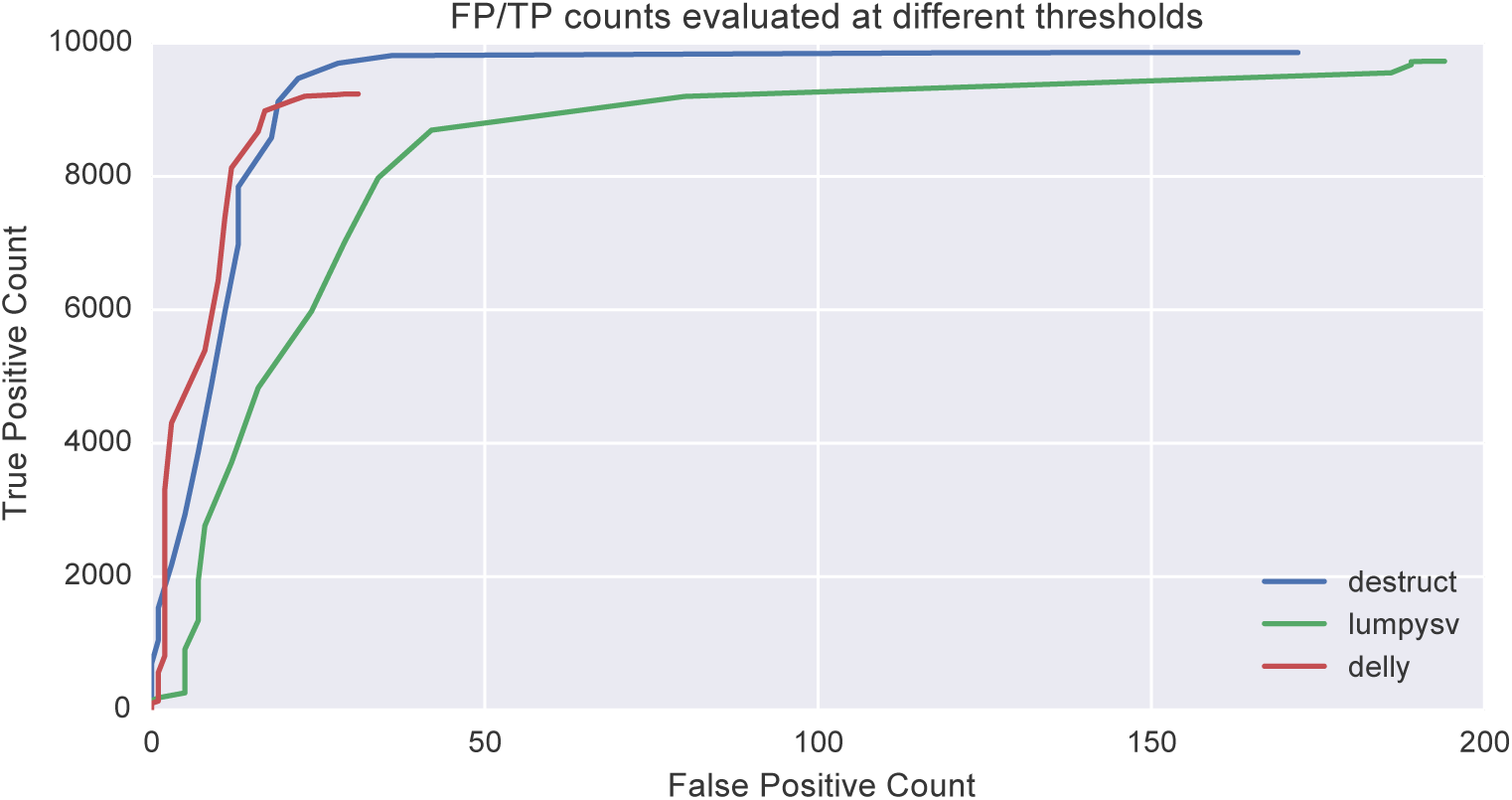
False positive versus true positive counts for a range of thresholds for destruct, delly, and lumpysv based on simulated breakpoints.

## 4 Availability

The deStruct source code including benchmarking scripts are available from http://bitbucket.org/dranew/destruct.

## References

[1] K. Chen, J. W. Wallis, M. D. McLellan, D. E. Larson, J. M. Kalicki, C. S. Pohl, S. D. McGrath, M. C. Wendl, Q. Zhang, D. P. Locke, X. Shi, R. S. Fulton, T. J. Ley, R. K. Wilson, L. Ding, and E. R. Mardis. Breakdancer: an algorithm for high-resolution mapping of genomic structural variation. Nat Methods, 6(9):677–81, Sep 2009.

[2] A. D. Ewing, K. E. Houlahan, Y. Hu, K. Ellrott, C. Caloian, T. N. Yamaguchi, J. C. Bare, C. P’ng, D. Waggott, V. Y. Sabelnykova, ICGC-TCGA DREAM Somatic Mutation Calling Challenge participants, M. R. Kellen, T. C. Norman, D. Haussler, S. H. Friend, G. Stolovitzky, A. A. Margolin, J. M. Stuart, and P. C. Boutros. Combining tumor genome simulation with crowdsourcing to benchmark somatic single-nucleotide-variant detection. Nat Methods, 12(7):623–30, Jul 2015.

[3] M. Griffith, O. L. Griffith, S. M. Smith, A. Ramu, M. B. Callaway, A. M. Brummett, M. J. Kiwala, A. C. Coffman, A. A. Regier, B. J. Oberkfell, G. E. Sanderson, T. P. Mooney, N. G. Nutter, E. A. Belter, F. Du, R. L. Long, T. E. Abbott, I. T. Ferguson, D. L. Morton, M. M. Burnett, J. V. Weible, J. B. Peck, A. Dukes, J. F. McMichael, J. T. Lolofie, B. R. Derickson, J. Hundal, Z. L. Skidmore, B. J. Ainscough, N. D. Dees, W. S. Schierding, C. Kandoth, K. H. Kim, C. Lu, C. C. Harris, N. Maher, C. A. Maher, V. J. Magrini, B. S. Abbott, K. Chen, E. Clark, I. Das, X. Fan, A. E. Hawkins, T. G. Hepler, T. N. Wylie, S. M. Leonard, W. E. Schroeder, X. Shi, L. K. Carmichael, M. R. Weil, R. W. Wohlstadter, G. Stiehr, M. D. McLellan, C. S. Pohl, C. A. Miller, D. C. Koboldt, J. R. Walker, J. M. Eldred, D. E. Larson, D. J. Dooling, L. Ding, E. R. Mardis, and R. K. Wilson. Genome modeling system: A knowledge management platform for genomics. PLoS Comput Biol, 11(7):e1004274, Jul 2015.

[4] F. Hormozdiari, C. Alkan, E. E. Eichler, and S. C. Sahinalp. Combinatorial algorithms for structural variation detection in high-throughput sequenced genomes. Genome Res, 19(7):1270–8, Jul 2009.

[5] B. Langmead, C. Trapnell, M. Pop, and S. L. Salzberg. Ultrafast and memory-efficient alignment of short dna sequences to the human genome. Genome Biol, 10(3):R25, 2009.

[6] R. M. Layer, C. Chiang, A. R. Quinlan, and I. M. Hall. Lumpy: a probabilistic framework for structural variant discovery. Genome Biol, 15(6):R84, Jun 2014.

[7] H. Li and R. Durbin. Fast and accurate short read alignment with burrows-wheeler transform. Bioinformatics, 25(14):1754–60, Jul 2009.

[8] H. Li and R. Durbin. Fast and accurate long-read alignment with burrows-wheeler transform. Bioinformatics, 26(5):589–95, Mar 2010.

[9] A. McPherson, A. Roth, C. Chauve, and S. C. Sahinalp. Joint inference of genome structure and content in heterogeneous tumor samples. In International Conference on Research in Computational Molecular Biology, pages 256–258. Springer, 2015.

[10] R. M. Neal. Markov chain sampling methods for dirichlet process mixture models. Journal of computational and graphical statistics, 9(2):249–265, 2000.

[11] T. Rausch, T. Zichner, A. Schlattl, A. M. Stütz, V. Benes, and J. O. Korbel. Delly: structural variant discovery by integrated paired-end and split-read analysis. Bioinformatics, 28(18):i333–i339, Sep 2012.

[12] A. M. S. Shrestha and M. C. Frith. An approximate bayesian approach for mapping paired-end dna reads to a reference genome. Bioinformatics, 29(8):965–72, Apr 2013.

